# An Autoantigen Profile from Jurkat T-Lymphoblasts Provides a Molecular Guide for Investigating Autoimmune Sequelae of COVID-19

**DOI:** 10.1101/2021.07.05.451199

**Authors:** Julia Y. Wang, Wei Zhang, Michael W. Roehrl, Victor B. Roehrl, Michael H. Roehrl

## Abstract

In order to understand autoimmune phenomena contributing to the pathophysiology of COVID-19 and post-COVID syndrome, we have been profiling autoantigens (autoAgs) from various cell types. Although cells share numerous autoAgs, each cell type gives rise to unique COVID-altered autoAg candidates, which may explain the wide range of symptoms experienced by patients with autoimmune sequelae of SARS-CoV-2 infection. Based on the unifying property of affinity between autoantigens (autoAgs) and the glycosaminoglycan dermatan sulfate (DS), this paper reports 140 candidate autoAgs identified from proteome extracts of human Jurkat T-cells, of which at least 105 (75%) are known targets of autoantibodies. Comparison with currently available multi-omic COVID-19 data shows that 125 (89%) of DS-affinity proteins are altered at protein and/or RNA levels in SARS-CoV-2-infected cells or patients, with at least 94 being known autoAgs in a wide spectrum of autoimmune diseases and cancer. Protein alterations by ubiquitination and phosphorylation in the viral infection are major contributors of autoAgs. The autoAg protein network is significantly associated with cellular response to stress, apoptosis, RNA metabolism, mRNA processing and translation, protein folding and processing, chromosome organization, cell cycle, and muscle contraction. The autoAgs include clusters of histones, CCT/TriC chaperonin, DNA replication licensing factors, proteasome and ribosome proteins, heat shock proteins, serine/arginine-rich splicing factors, 14-3-3 proteins, and cytoskeletal proteins. AutoAgs such as LCP1 and NACA that are altered in the T cells of COVID patients may provide insight into T-cell responses in the viral infection and merit further study. The autoantigen-ome from this study contributes to a comprehensive molecular map for investigating acute, subacute, and chronic autoimmune disorders caused by SARS-CoV-2.

## Introduction

The COVID-19 pandemic has been devastating. After initial recovery from acute SARS-CoV-2 infection, many people continue to suffer from lingering health problems (so called “long COVID” or post-COVID syndrome), such as fatigue, shortness of breath, joint pain, chest pain, muscle pain, loss of smell or taste, and other neurological problems. Although the underlying causes are far from clear, autoimmune effects are likely important contributors to chronic post-COVID disorders. To understand how SARS-CoV-2 infection may induce autoimmune responses, we are establishing a comprehensive COVID autoantigen atlas, i.e., all possible endogenous autoantigens (autoAgs) that may be rendered immunogenic by the viral infection. Because different tissues or cells may give rise to distinct pools of autoAgs, we have been profiling autoAgs from multiple human tissues and cell types, including human lung fibroblast HFL1 cells, human lung epithelial-like A549 cells, and B-lymphoblast HS-Sultan cells [1–4]. In this study, we report an autoantigen-ome identified from human Jurkat T-lymphoblast cells.

Our autoAg discovery is based a unifying mechanism of autoantigenicity that we have uncovered [5–8]. AutoAgs are the targets of autoantibodies (autoAbs) and T-cell autoimmune responses. Typically, self-molecules are naturally tolerated by the immune system and do not provoke autoimmune responses. However, certain self-molecules transform into autoAgs and become targets of autoimmune attacks. Thus far, hundreds of autoAgs with seemingly no obvious structural or functional commonality have been identified across various autoimmune diseases and cancers. Our studies have demonstrated that autoAgs do, in fact, share common properties. AutoAgs are commonly released by apoptotic cells, and we found that the glycosaminoglycan dermatan sulfate (DS) has peculiar affinity to apoptotic cells and their autoAgs [5, 7, 8]. DS and autoAgs can form affinity complexes and cooperatively stimulate autoreactive B1 cells and autoantibody production [5, 7, 8]. Based on autoAg-DS affinity, we have identified several hundred autoAgs from various cells and tissues [1–3, 9–11].

A variety of autoantibodies have been identified in COVID-19 patients [12–22]. Children infected with SARS-CoV-2 who develop the rare multisystem inflammatory syndrome show multiple autoAbs, including classical antinuclear antigen (ANA) autoAbs and specific autoAbs recognizing endothelial, gastrointestinal, or immune cell autoAgs [12, 13]. ANA autoAbs are also frequently detected in COVID-19 patients with acute respiratory syndrome or other critical conditions [14–16] and in COVID patients with no previous clinical record of autoimmune diseases [17]. A high frequency of cerebrospinal fluid autoAbs is found in COVID patients with neurological symptoms [18]. New-onset autoAbs were detected in a significant proportion of hospitalized COVID-19 patients and were positively correlated with immune responses to SARS-CoV-2 proteins [20]. Overall, an increasing number of observations suggest a positive correlation between emergence of autoAbs and an adverse clinical course of COVID-19.

As revealed by our prior studies, SARS-CoV-2 infection may induce numerous molecular changes in the host and transform naturally non-antigenic self-molecules to antigenic autoAgs [1–3]. In order to better understand the possible extent of autoimmune disorders caused by SARS-CoV-2, we are building a comprehensive catalog of all possible intrinsic autoAgs across cell and tissue types related to the viral infection. Herein, we report a profile of autoAgs identified from human Jurkat T-cells using our DS-affinity enrichment approach, which will provide valuable molecular targets for understanding the diverse autoimmune sequelae of COVID-19.

## Results and Discussion

### Autoantigen-ome of Jurkat cells identified by DS-affinity

Total proteins were extracted from Jurkat T-cells and fractionated in a DS-Sepharose affinity column. Proteins with increasing DS-affinity were eluted from the column with increasing ionic strength of salt. Fractions eluted with 0.4, 0.6, and 1.0 M NaCl correspond to proteins with intermediate, strong, and very strong DS-affinity, respectively. Mass spectrometry sequencing identified a total of 140 proteins from these three DS-affinity fractions (Table 1). The majority of proteins (120/140) were eluted with 0.4 M NaCl, 31 proteins were found in the 0.6 M NaCl elution, and 11 proteins were identified in the 1.0 M NaCl elution. Three proteins were detected redundantly in all three fractions (HIST4H4, H2AC1, RPLP2), 1H2BC1 was detected in both 0.6 and 1.0 M fractions, C1QBP was detected in both 0.4 and 1.0 M NaCl fractions, and 13 proteins were detected in both 0.4 and 0.6 M fractions.

**Table 1.**
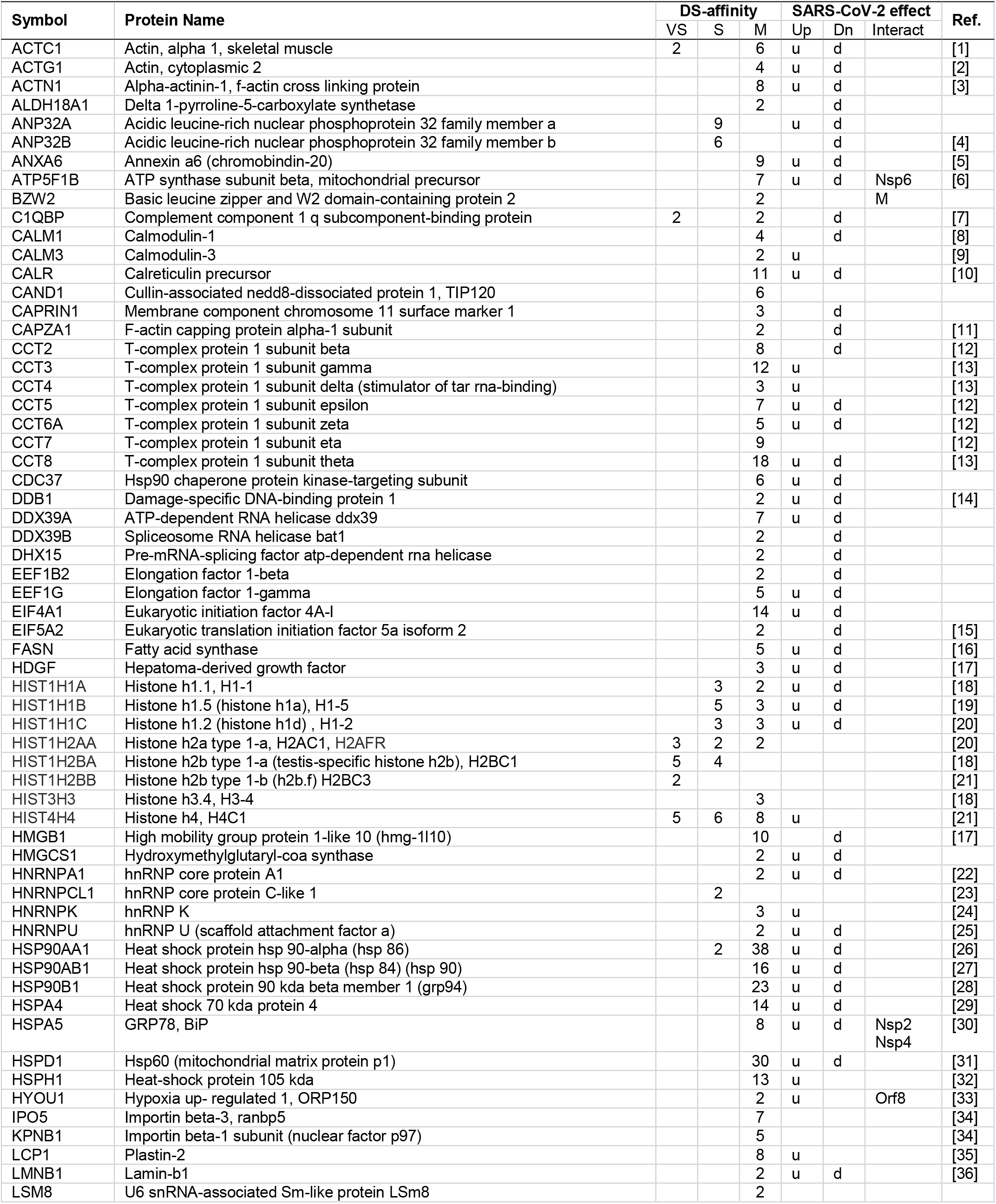
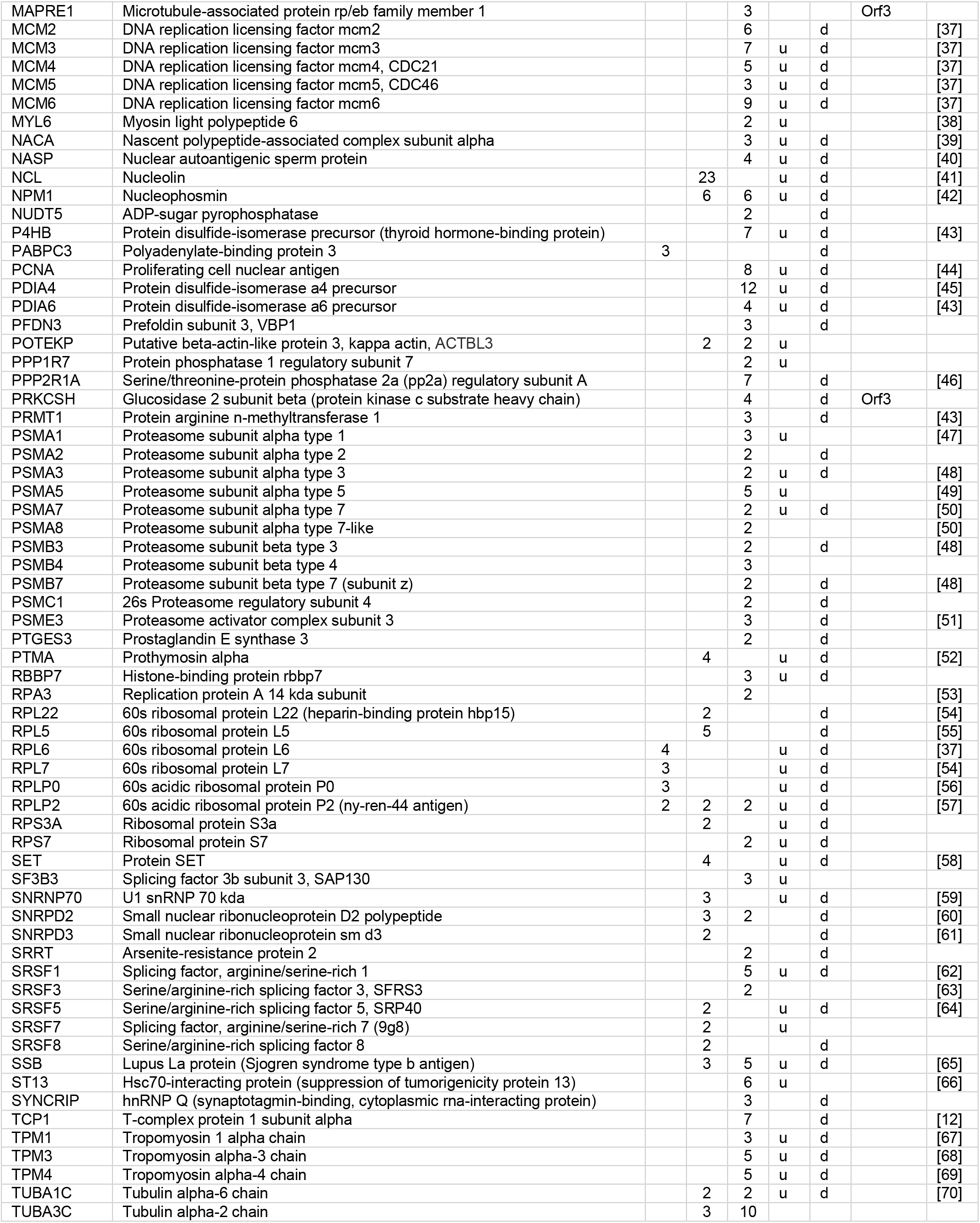
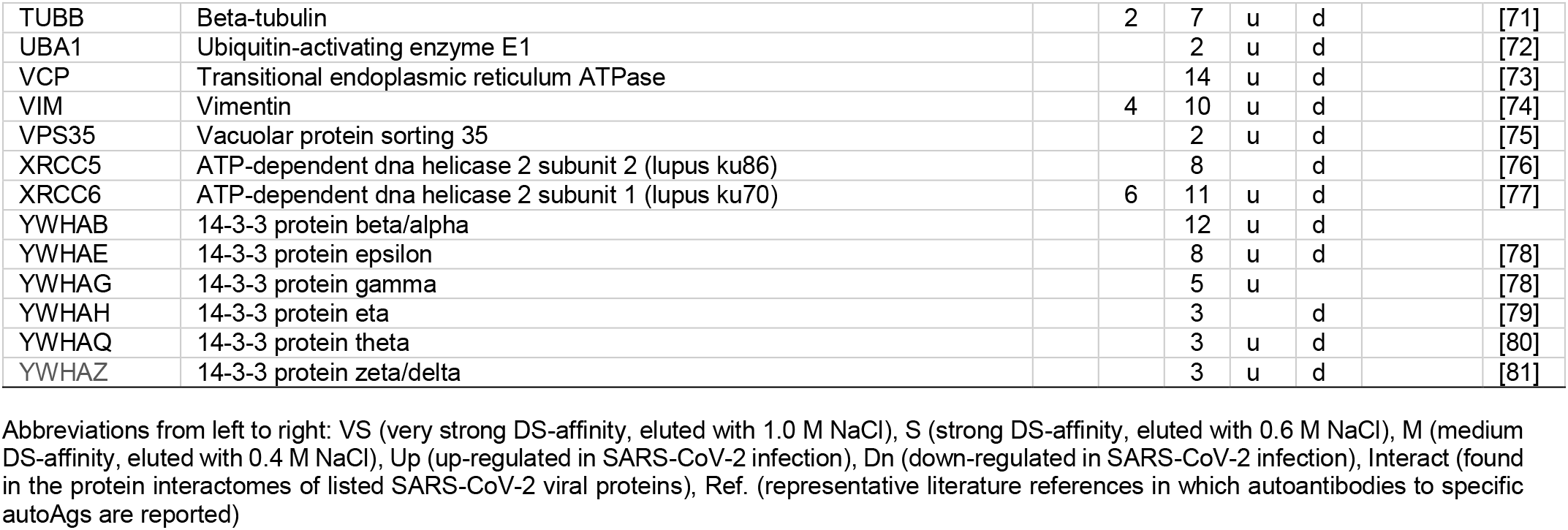
DS-affinity autoantigens from Jurkat T-cells and their alterations in SARS-CoV-2 infection

Remarkably, among the 140 DS-affinity proteins identified from Jurkat T-cells, at least 105 (75%) are known autoAgs, i.e., the existence of specific autoantibodies against these proteins has been reported in the literature (see references in Table 1). These autoantibody/autoAg pairs are found in a wide spectrum of autoimmune diseases as well as a variety of cancers. Although 36 of the DS-affinity proteins have not yet been reported as autoAgs, we suspect that most, if not all, are putative autoAgs awaiting serological confirmation. For example, 6 serine/arginine-rich splicing factors were identified by DS-affinity, but only 3 of them (SRSF1, SRSF3, SRSF5) have thus far been individually reported as autoAgs (Table 1). A serine/arginine-rich repeating octapeptide of Arg-Ser-Arg-Ser-Arg(Lys)-Glu(Asp)-Arg-Lys(Arg) has been found in several nuclear autoAgs such as U2AF 35- and 65-kDa splicing factors and 70-KDa U1 snRNP [23], and many other splicing factors have been reported as autoAgs, such as SF3B1 and SRSF2. Therefore, we suspect that the other 3 splicing factors (SRSF3B3, SRSF7, and SRSF8) identified by DS-affinity in this study are likely true autoAgs that are yet to be confirmed.

Proteins eluted with 1.0 M NaCl possess the strongest DS-affinity and, strikingly, 10/13 (90.9%) are known autoAgs (Table 1), indicating that increasing affinity to DS increases the propensity of a protein to be an autoAg, consistent with our prior findings [1–3, 5, 6, 9–11]. These include histones (H4, H2B types 1-a and 1-b, H2A type 1-a), 60S ribosomal proteins (P0, P2, L6, L7), ACTC1 (skeletal muscle actin), and C1QBP, and PABPC3 (polyadenylate-binding protein 3). Histones and ribosomal P proteins are hallmark autoAgs used in routine clinical tests of autoimmune diseases. Histone autoAbs are nearly always present in drug-induced systemic lupus erythematosus, and ribosomal P autoantibodies are tested for to aid in the differential diagnosis of lupus patients with neuropsychiatric symptoms. C1QBP has been repeatedly identified as a putative autoAg in several of our prior studies [1, 2, 9, 10] and was recently confirmed as an autoAg in the neurodegenerative disorder primary open-angle glaucoma [24]. Poly(A)-binding proteins bind the poly(A) tail of messenger RNAs and control mRNA stability and translation initiation. Although PABPC3 has not yet been reported as an autoAg, its paralog PABPC1 has been found to be an autoAg [25].

Proteins eluted with 0.6 M NaCl possess strong DS-affinity and, remarkably, 26/31 (83.9%) are known autoAgs (Table 1). Several well-known autoAgs are identified in this strong DS-affinity fraction, including 6 histone autoAgs, SSB (lupus La autoAg), XRCC6 (lupus Ku70 autoAg), 3 snRNP autoAgs (Sm D2, Sm D3, U1 70kD). Other autoAgs identified with strong DS-affinity include ANP32B, nucleolin, nucleophosmin, SET, HNRNPCL1, HSP90AA1, 3 ribosomal proteins (L22, L5, S3a), 3 serine/arginine-rich splicing factors, 3 tropomyosin subunits, prothymosin alpha, 3 tubulin subunits, vimentin, and T-complex protein 1 alpha. A few have not yet been confirmed as autoAgs, including ANP32A, kappa actin, and ribosomal protein 3A.

The 140 candidate autoAgs identified from Jurkat T-cells are not a random collection but are highly enriched in a few groups of proteins. Among them, there are 11 proteasomal proteins, 8 ribosomal proteins, 8 histones, 8 T-complex protein (CCT/TriC) subunits, 7 heat shock proteins, 6 splicing factors, 6 14-3-3 proteins, 5 DNA replication licensing factors (or minichromosome maintenance proteins), 5 DNA or RNA helicases, and 4 hnRNPs.

Protein-protein interaction network analysis by STRING [26] reveals that the DS-affinity autoantigen-ome is highly connected (Fig. 1). There are 787 interactions at high confidence level (vs. 284 expected; enrichment p-value <1.0e-16). These DS-affinity proteins are enriched in several clusters and significantly associated with the cell cycle, protein folding, chromosome organization, RNA splicing, translation, and muscle contraction (Fig. 1). There are 36 DS-affinity proteins associated with the cell cycle, particularly the G2/M checkpoints (26 proteins), the G2/M DNA damage checkpoint, and the G1/S and G2/M transitions.

**Fig. 1.**
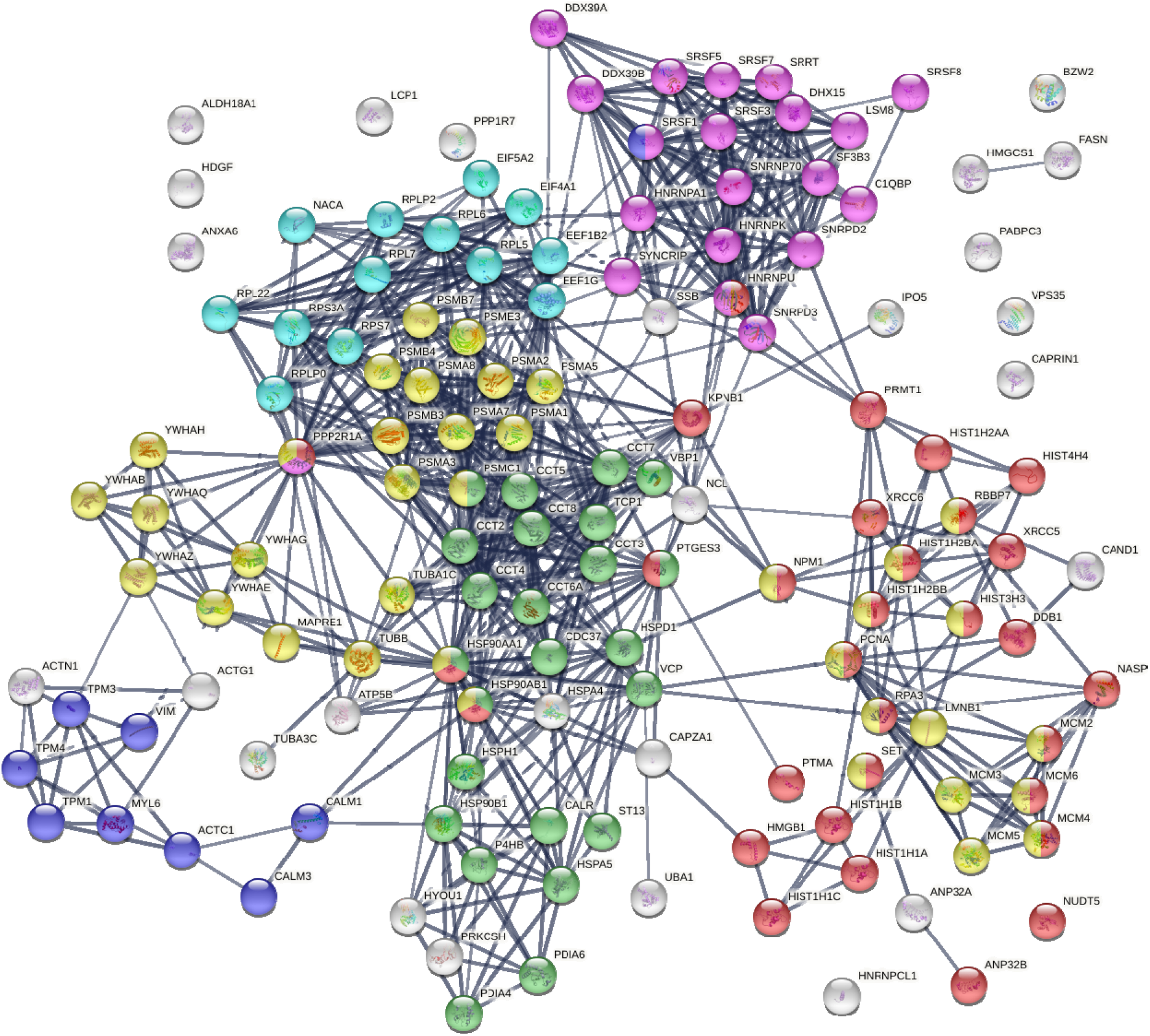
The autoantigen-ome from Jurkat T-cells identified by DS affinity. Lines represent protein-protein interactions at high confidence levels. Marked proteins are associated with cell cycle (37 proteins, yellow), chromosome organization (31 proteins, red), RNA splicing (20 proteins, pink), translation (13 proteins, aqua), protein folding (24 proteins, green), and muscle contraction (9 proteins, blue).

Pathway and process enrichment analyses by Metascape [27] also reveal that proteins of the DS-affinity autoantigen-ome are significantly associated with cellular response to stress, protein folding, and protein localization to organelles (Fig. 2A). In addition, they are associated with kinase maturation complex 1, spliceosome, HSF1 activation (activates gene expression in response to a variety of stresses), protein processing in the endoplasmic reticulum, VEGFA-VEGFR2 signaling (major pathway that activates angiogenesis), apoptosis-induced DNA fragmentation, and 17S U2 snRNP.

**Fig. 2.**
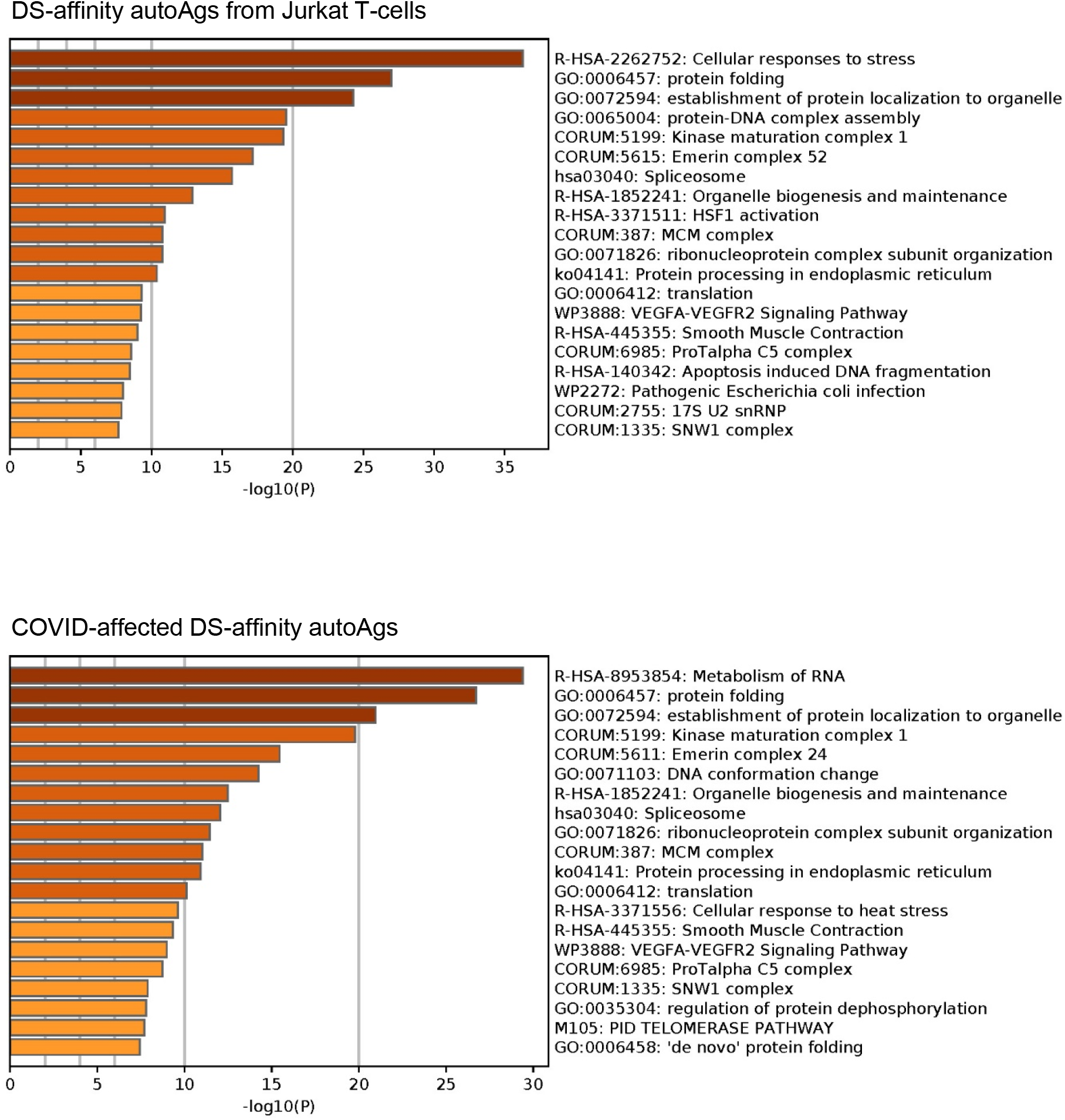
Top 20 enriched pathways and processes among COVID-altered DS-affinity proteins. Top chart: 140 proteins identified by DS-affinity from Jurkat T-cells. Bottom chart: 125 DS-affinity proteins that are altered in SARS-CoV-2 infection.

### DS-affinity autoantigen-ome related to COVID-19

To find out how many of the DS-affinity autoAgs identified from Jurkat T-cells are affected by in SARS-CoV-2 infection, we searched for them in a multi-omic COVID database compiled by Coronascape [27–47]. Among the 140 DS-affinity proteins identified in our study, 125 (89.3%) are affected by SARS-CoV-2 infection, and at least 94 (of the 125; 75.2%) are known autoAgs (Table 1 and Supplemental Table 1). Among the COVID-altered DS-affinity proteins, 17 are up-regulated only, 35 are down-regulated only, and 71 are altered (up or down depending on study conditions) at protein and/or RNA levels in SARS-CoV-2 infected cells. The COVID database was assembled from different cell and patient tissue types by multiple research laboratories using different technologies, including proteomics, phosphoproteomics, ubiquitinomics, and bulk and single-cell RNA sequencing.

Six DS-affinity proteins are found in the interactomes of SARS-CoV-2 viral proteins, i.e., these host proteins interact directly or indirectly with the viral proteins [29, 40, 44]. Specifically, HSPA5 (GRP78/BiP) interacts with Nsp2 and Nsp4, HYOU1 interacts with Orf8, PRKCSH and MAPRE1 interact with Orf3, and BZW2 interacts with the viral M protein. HSPA5/BiP (binding immunoglobulin protein) has been consistently identified by DS-affinity in our previous studies, and we have also recently reported that DS-BiP association plays important roles in regulating precursor autoreactive B1 cells [8]. HYOU1 (hypoxia up-regulated protein 1) was also found overexpressed at protein level in the urine of COVID-19 patients and up-regulated at mRNA level in B cells from 4 patients out of a cohort of 7 hospitalized COVID-19 patients [33, 48]. HYOU1 belongs to the heat shock protein 70 family, accumulates in the endoplasmic reticulum under hypoxic conditions, and has been shown to be up-regulated in tumors. PRKCSH (glucosidase 2 subunit beta) is an N-linked glycan processing enzyme in the endoplasmic reticulum, and mutations of this gene have been associated with the autosomal dominant polycystic liver disease. MAPRE1 (microtubule-associated protein RP/EB family member 1) binds the plus-end of microtubules and regulates microtubule cytoskeleton dynamics. BZW2 (basic leucine zipper and W2 domain 2) may be involved in neuronal differentiation and is associated with congenital hypomyelinating neuropathy.

Similar to the 140 DS-affinity protein autoantigen-ome, the 125 COVID-altered DS-affinity proteins are most significantly associated with RNA metabolism and protein folding (Fig. 2B). In addition, they are associated with establishment of protein localization to organelles, kinase maturation complex 1, emerin complex 24, DNA conformation change, spliceosome, cellular response to heat stress, smooth muscle contraction, VEGFA-VESFR2 signaling pathway, prothymosin alpha C5 complex, regulation of protein dephosphorylation, and telomerase pathway (Fig. 2B). Protein-protein interaction network analysis also confirms that the COVID-altered DS-affinity protein network is strongly associated with mRNA processing, translation, chromosome organization, protein processing in the endoplasmic reticulum, CCT/TriC chaperonin, and apoptosis (Fig. 3).

**Fig. 3.**
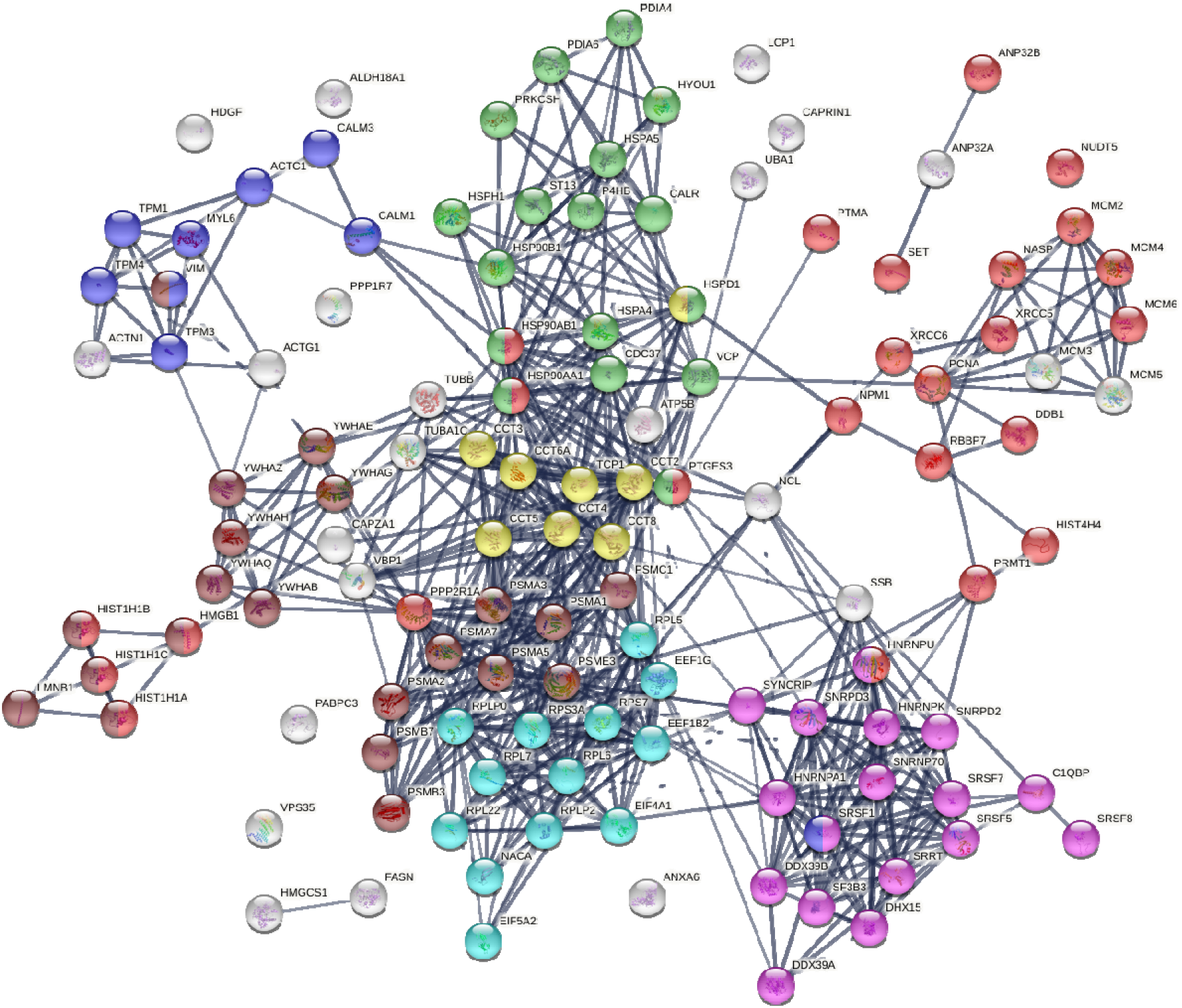
DS-affinity proteins that are altered by SARS-CoV-2 infection. Lines represent protein-protein interactions at high confidence levels. Marked proteins are associated with chromosome organization (25 proteins, red), mRNA processing (17 proteins, pink), translation (13 proteins, aqua), protein processing in endoplasmic reticulum (green, 17 proteins), muscle contraction (9 proteins, blue), TCP-1/cpn60 chaperonin (yellow, 8 proteins), and apoptosis (21 proteins, brown).

Nine COVID-altered DS-affinity proteins are associated with muscle contraction, including ACTC1, CALM1, CALM3, MYL6, TPM1, TPM3, TPM4, SRSF1, and VIM. All of these proteins are known autoAgs (Table 1). CALM1 has recently been identified as one of the autoAgs in multisystem inflammatory syndrome in children from SARS-CoV-2 infection [13]. Six 14-3-3 proteins are identified, all of which are autoAgs. Presence of 14-3-3 proteins in cerebrospinal fluid, a marker of ongoing neurodegeneration, has been detected in COVID-19 patients [49].

### AutoAgs from altered phosphorylation and ubiquitination

Thirty-eight of the 125 COVID-affected DS-affinity proteins have phosphorylation changes in SARS-CoV-2 infection (Fig. 4). Their molecular functions include histone binding (6 proteins), RNA binding (10 proteins), helicase activity (5 proteins), ATP binding (12 proteins), DNA binding (14 proteins), and hydrolase activity (11 proteins). These COVID-altered phosphoproteins are significantly associated with gene expression, chromosome organization, and mRNA metabolism. Chromosome-associated proteins are particularly related to DNA conformation change (XRCC6, SET, NPM1, HIST1H1C, HIST1H1B, RBBP7, NASP, MCMs) and DNA replication (MCM2, MCM3, MCM4, NASP, RBBP7, and SET). mRNA-associated proteins are related to mRNA splicing (SRSF1, SRSF7, SRRT, HNRNPA1, HNRPNK, HNRNPU, DDX39A) and RNA 3’-end processing (DDX39A, SRSF7, SRSF1, SSB). In addition, nuclear matrix protein lamin-B1, nucleolar protein nucleolin, vacuolar protein sorting-associated protein VPS35, vimentin, fatty acid synthetase FASN, protein phosphatase 1 regulatory subunit PPP1R7, and HDGF (hepatoma-derived growth factor) are altered by phosphorylation.

**Fig. 4.**
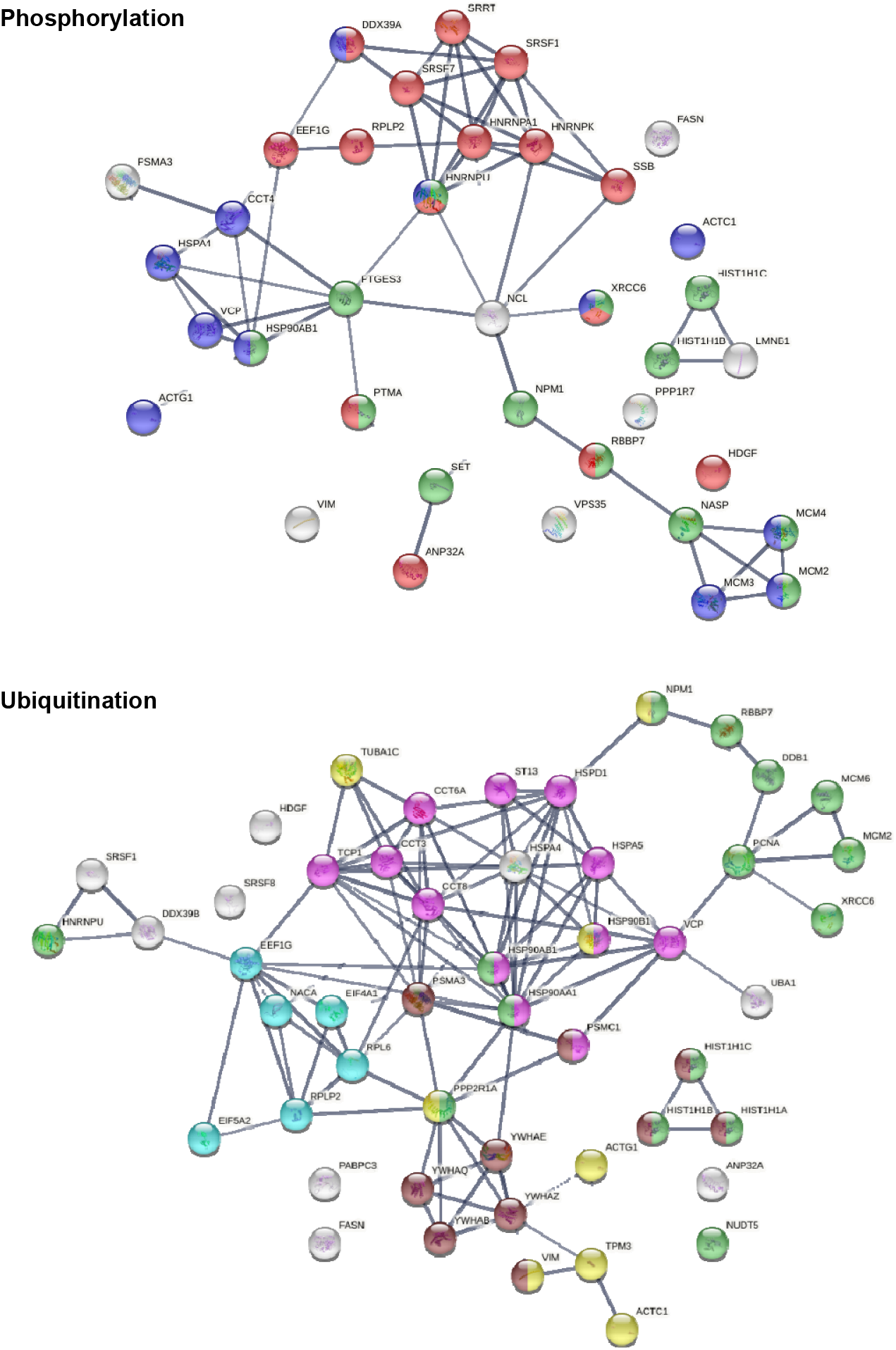
DS-affinity proteins that show changes in phosphorylation or ubiquitination in SARS-CoV-2 infection. Phosphorylation: marked proteins are associated with gene expression (15 proteins, red), chromosome organization (13 proteins, green), and ATP binding (12 proteins, blue). Ubiquitination: marked proteins are associated with protein folding (12 proteins, pink), chromosome organization (15 protein, green), translation (6 protein, aqua), cytoskeleton (8 proteins, yellow), and apoptosis (10 proteins, brown).

Among the 125 COVID-affected DS-affinity proteins, 50 are altered by ubiquitination in SARS-CoV-2 infection (Fig. 4). These proteins are associated with apoptosis, chromosome organization, protein folding, translation, cell cycle, and cytoskeleton. Proteins related to apoptosis include linker histones (HIST1H1A, HIST1H1B, HIST1H1C), 14-3-3 proteins (YWHAB, YWHAE, YWHAQ, YWHAZ), and proteasome proteins (PSMA3, PSMC1). Proteins related to the cell cycle include PNCA, MCM2, MCM6, and 14-3-3 proteins. Five heat shock proteins and 4 subunits of chaperonin CCT/TriC are ubiquitinated. Other interesting ubiquitinated proteins include NACA (nascent polypeptide-associated complex subunit alpha), DDB1 (DNA damage-binding protein 1), NUDT5 (ADP-sugar pyrophosphatase), and NPM1 (nucleophosmin). Ubiquitination is typically the “kiss of death” modification that marks proteins destined for degradation by the proteasome, although ubiquitination may also modulate protein interaction and activity. Intriguingly, we identified UBA1 (ubiquitin-like modifier-activating enzyme 1), which catalyzes the first step in ubiquitin conjugation and plays a central role in ubiquitination, as a ubiquitination-altered DS-affinity autoAg, which is consistent with our previous studies [1, 2].

### DS-affinity proteins altered in T cells of COVID-19 patients

Because Jurkat cells were established from human T-cell lymphoblastic leukemia, we searched for DS-affinity proteins that were altered in T cells of seven COVID-19 patients [33]. Five proteins (LCP1, CALR, HSPA5, HSP90AA1, HSP90AB1) were up-regulated in CD4+ T cells, and 13 proteins (LCP1, CALR, HSPA5, HSP90AA1, HSP90AB1, HSPD1, HSPH1, MCM4, VIM, PTMA, TUBB, H1-2, LMNB1) were up-regulated in CD8+ T cells of COVID-19 patients. Three proteins (ACTG1, EEF1B2, SRSF5) were down-regulated in the CD4+ T cells, and 3 proteins (ATCG1, EEF1B2, NACA) were down-regulated in CD8+ T cells. Remarkably, all up-regulated DS-affinity proteins are known autoAgs (Table 1). NACA, ACTG1, and SRSF5, which were down-regulated at the mRNA level, are also known autoAgs. EEF1B2 (or EEF1B, elongation factor 1-beta) has not been identified as an autoAg, although other similar elongation factors such as EEF1A and EF2 are known autoAgs (see references in Table 1).

Among up-regulated proteins, LCP1 was up-regulated in CD4+ T cells of 2 patients (out of 4 patients with available data) and in CD8+ T cells of 2 patients (out of 5 patients with available data), with one of the patients having LCP1 up in both CD4+ and CD8+ T cells. Up-regulation of heat shock proteins, particularly HSPA5 and HSP90AA1, was detected in CD4+ T cells of 2 patients and CD8+ T cells of 1 patient. MCM4 up-regulation was detected in CD8+ T cells of 3 out of 6 patients. Among down-regulated proteins, NACA was detected in CD4+ T cells of 1 patient and CD8+ T cells of all 3 patients whose data were available. EEF1B2 was down in CD4+ T cells of 3 patients (out of 5 with available data) and down in CD8+ T cells of 2 out of 3 patients. ACTG1 down-regulation was detected in CD4+ T cells of 2 patients and CD8+ T cells of 1 patient. SRSF5 was down in CD4+ T cells of 3 out of 5 patients.

Among these T-cell-altered proteins, LCP1 and NACA are perhaps most interesting. LCP1 (plastin-2, an actin binding protein) has been found to play a significant role in T cell activation in response to co-stimulation through TCR/CD3 and regulates the stability of the immune synapse of naïve and effector T cells [50]. NACA (nascent polypeptide-associated complex subunit alpha) binds to newly synthesized polypeptide chains as they emerge from the ribosome, blocks their interaction with the signal recognition particle, and prevents inappropriate targeting of non-secretory polypeptides to the endoplasmic reticulum. NACA is an IgE autoAg in atopic dermatitis patients with chronic skin manifestations [51]. The significance of these T-cell proteins in COVID-19 and autoimmunity merits further study.

## Conclusion

In order to establish a comprehensive COVID-19 autoantigen-ome, we have been profiling autoAgs from different cell and tissue types. Compared to other cells we have examined, Jurkat T-cells contain relatively fewer DS-affinity autoAgs than HFL1 lung fibroblasts, A549 lung epithelial cells, HS-Sultan B-lymphoblasts, and HEp-2 fibroblasts. Although cells share numerous autoAgs, each cell type gives rise to unique COVID-altered autoAg candidates, which may explain the wide range of symptoms experienced by patients with autoimmune sequelae of SARS-CoV-2 infection. We believe that our effort of discovering autoAgs across different cell types provides a comprehensive and valuable autoAg database for better understanding of autoimmune diseases and post-COVID-19 health problems.

## Materials and Methods

### Jurkat T-cell culture

The human T lymphoblast Jurkat cell line was obtained from the ATCC (Manassas, VA) and cultured in complete RPMI-1640 medium. The growth medium was supplemented with 10% fetal bovine serum and a penicillin-streptomycin-glutamine mixture (Thermo Fisher). The cells were grown at 37 °C in a CO_2_ incubator.

### Protein extraction

Protein extraction was performed as previously described [6]. In brief, Jurkat cells were lysed with 50 mM phosphate buffer (pH 7.4) containing the Roche Complete Mini protease inhibitor cocktail and then homogenized on ice with a microprobe sonicator until the turbid mixture turned nearly clear with no visible cells left. The homogenate was centrifuged at 10,000 g at 4 °C for 20 min, and the total protein extract in the supernatant was collected. Protein concentration was measured by absorbance at 280 nm using a NanoDrop UV-Vis spectrometer (Thermo Fisher).

### DS-Sepharose resin preparation

The DS-affinity resins were synthesized as previously described [6, 9]. In brief, 20 ml of EAH Sepharose 4B resins (GE Healthcare Life Sciences) were washed with distilled water three times and mixed with 100 mg of DS (Sigma-Aldrich) in 10 ml of 0.1 M MES buffer, pH 5.0. About 100 mg of N-(3-dimethylaminopropyl)-N’-ethylcarbodiimide hydrochloride (Sigma-Aldrich) powder was added, and another 100 mg was added after 8 h of reaction. The reaction proceeded by mixing on a rocker at 25 °C for 16 h. The coupled resins were washed with water and equilibrated with 0.5 M NaCl in 0.1 M acetate (pH 5.0) and 0.5 M NaCl in 0.1 M Tris (pH 8.0).

### DS-affinity fractionation

The total proteomes extracted from Jurkat cells were fractionated in a DS-Sepharose column [6]. About 40 mg of proteins in 40 ml of 10 mM phosphate buffer (pH 7.4; buffer A) were loaded onto the DS-affinity column at a rate of 1 ml/min. Unbound and weakly bound proteins were removed with 60 ml of buffer A and then 40 ml of 0.2 M NaCl in buffer A. The remaining bound proteins were eluted in step gradients of 40 ml each of 0.4 M, 0.6 M, and 1.0 M NaCl in buffer A. Fractions were desalted and concentrated with 5-kDa cut-off Vivaspin centrifugal filters (Sartorius). Fractionated proteins were separated in 1-D SDS-PAGE in 4-12% Bis-Tris gels, and each gel lane was divided into two or three sections for sequencing.

### Mass spectrometry sequencing

Protein sequencing was performed at the Taplin Biological Mass Spectrometry Facility at Harvard Medical School. Proteins in gels were digested with sequencing-grade trypsin (Promega) at 4 °C for 45 min. Tryptic peptides were separated in a nano-scale C18 HPLC capillary column and analyzed in an LTQ linear ion-trap mass spectrometer (Thermo Fisher). Peptide sequences and protein identities were assigned by matching the measured fragmentation pattern with proteins or translated nucleotide databases using Sequest. All data were manually inspected. Proteins with ≥2 peptide matches were considered positively identified.

### COVID data comparison

DS-affinity proteins were compared with currently available COVID-19 multi-omic data compiled in the Coronascape database (as of 02/22/2021) [27–48]. These data have been obtained with proteomics, phosphoproteomics, interactome, ubiquitome, and RNA-seq techniques. Up- and down-regulated proteins or gene transcripts were identified by comparing cells infected vs. uninfected by SARS-CoV-2 or COVID-19 patients vs. healthy controls. Similarity searches were conducted to identify DS-affinity proteins that are up- and/or down-regulated in viral infection at any omic level.

### Protein network analysis

Protein-protein interactions were analyzed by STRING [26]. Interactions include both direct physical interaction and indirect functional associations, which are derived from genomic context predictions, high-throughput lab experiments, co-expression, automated text mining, and previous knowledge in databases. Each interaction is annotated with a confidence score from 0 to 1, with 1 being the highest, indicating the likelihood of an interaction to be true. Pathways and processes enrichment were analyzed with Metascape [27], which utilizes various ontology sources such as KEGG Pathway, GO Biological Process, Reactome Gene Sets, Canonical Pathways, CORUM, TRRUST, and DiGenBase. All genes in the genome were used as the enrichment background. Terms with a p-value <0.01, a minimum count of 3, and an enrichment factor (ratio between the observed counts and the counts expected by chance) >1.5 were collected and grouped into clusters based on their membership similarities. The most statistically significant term within a cluster was chosen to represent the cluster.

### Autoantigen literature text mining

Every DS-affinity protein identified in this study was searched for specific autoantibodies reported in the PubMed literature. Search keywords included the MeSH keyword “autoantibodies”, the protein name or its gene symbol, or alternative names and symbols. Only proteins for which specific autoantibodies are reported in PubMed-listed journal articles were considered “confirmed” or “known” autoAgs in this study.

## Acknowledgements

We are grateful to Dr. Jung-hyun Rho for his technical assistance to the study. We thank Ross Tomaino and the Taplin Mass Spectrometry Facility of Harvard Medical School for expert service with protein sequencing.

## Authors’ contributions

JYW conducted the study and wrote the manuscript. WZ performed some of the experiments. MWR and VBR assisted with the study and manuscript preparation. MHR consulted on the study and edited the manuscript. All authors have approved the manuscript.

